# BioNAR: An Integrated Biological Network Analysis Package in Bioconductor

**DOI:** 10.1101/2023.02.08.527636

**Authors:** Colin McLean, Anatoly Sorokin, T. Ian Simpson, J Douglas Armstrong, Oksana Sorokina

**Affiliations:** School of informatics, University of Edinburgh, UK; Biological Systems Unit, Okinawa Institute of Science and Technology, Okinawa, Japan; Computational Biomedicine Institute (IAS-5 / INM-9), Forschungszentrum Jülich, Jülich, Germany; Health Economics and Data Science, Institute for Genetics and Cancer, University of Edinburgh, UK

## Abstract

Biological function in protein complexes emerges from more than just the sum of their parts: Molecules interact in a range of different subcomplexes and transfer signals/information around internal pathways. Modern proteomic techniques are excellent at producing a parts-list for such complexes, but more detailed analysis demands a network approach linking the molecules together and analyzing the emergent architectural properties. Methods developed for the analysis of networks in social sciences have proven very useful for splitting biological networks into communities leading to the discovery of sub-complexes enriched with molecules associated with specific diseases or molecular functions that are not apparent from the constituent components alone. Here we present the Bioconductor package BioNAR which supports step-by-step analysis of biological/biomedical networks with the aim of quantifying and ranking each of the network’s vertices based on network topology and clustering. Examples demonstrate that while BioNAR is not restricted to proteomic networks, it can predict a protein’s impact within multiple complexes, and enables estimation of the co-occurrence of meta-data, i.e., diseases and functions across the network, identifying the clusters whose components are likely to share common function and mechanisms. The package is available from Bioconductor release 3.16: https://bioconductor.org/packages/release/bioc/html/BioNAR.html

**Author Biographies:** Colin McLean holds a PhD in Experimental Particle Physics from the University of Edinburgh. He is currently a Senior Research Fellow in Health Economics and Data Science at the Institute for Genetics and Cancer at the University of Edinburgh. His research interests include applied network and data science in the biomedical domain.

Anatoly Sorokin holds PhD Degree in Biophysics and is a Senior computational biologist in the Biological Systems Unit, Okinawa Institute of Science and Technology. His research interests include graph-based analysis, constraint-base, dynamics and rule-based modelling and application areas include systems biology, bioinformatics and microbiomics. Ian Simpson has a DPhil in Genetics (Oxford 2000) and is currently Director of the UKRI Centre for Doctoral Training in Biomedical Artificial Intelligence and a Reader in Biomedical Informatics in the School of Informatics at The University of Edinburgh. His research interests lie at the boundary between Informatics and Biomedicine and focus on jointly modelling molecular and clinical data to improve our understanding of genetic disease.

J Douglas Armstrong holds a PhD in Molecular Genetics (Glasgow 1995) and is currently Professor of Systems Neurobiology at the School of Informatics at Edinburgh University. His research interests focus on structure/function mapping in the brains of model organisms. Oksana Sorokina holds a PhD in Systems Biology (Edinburgh 2010) and is a Senior Researcher at the School of Informatics at Edinburgh University. Her expertise is in the computational analysis of complex datasets primarily proteomics and the integration of genetic and other omic data types to understand molecular complexes at the systems biology level.

## Introduction

Biotechnology has made rapid advances in recent years with massive steps forward in both the sensitivity and throughput of methods to analyse biological samples across multiple levels. The most widely known advances are those in the high-throughput sequencing of DNA and RNA but there have been step changes in a range of areas including proteomics, metabolomics, and connectomics. These data are gathered for a purpose. The vast majority of biological processes that underpin our health and well-being both internally and at our social and environmental levels are really quite poorly understood beyond the role of a limited set of specific components. Integrating these components into a network model allows the identification of key functional interactions, pathways and complexes which are required for the healthy function and that, when perturbed, lead to disease.

Communication, or information transfer, between the components of the network is a common feature across biological scales. Therefore, it is not entirely surprising that methods designed to analyse information flow or communication in other networks have proven to be well suited to the analysis of biological networks. The history of algorithm development for the analysis of social networks has proven to be a rich hunting ground for useful methods in molecular biology, where many of these methods have been applied to the analysis of molecular interaction networks [1-3].

Proteomic data are typically represented via static undirected protein-protein interaction (PPI) networks, where vertices represent the proteins obtained from mass-spectrometry experiments and edges represent the structural protein interactions connecting them. From a PPI network one can extract many statistical measures of the network’s topology and its fundamental properties and use these to gain insight and make predictions about the underlying data [4, 5]. Centrality measures such as vertex degree and betweenness, can help reveal important or influential proteins within the network (see Case Studies). ‘Scale-free’ [6] properties and small world paths found in many biological networks are used to identify ‘hub’ molecules, which often encode disease related proteins [4]. Another useful network measure is graph entropy, which can provide an estimate of a protein’s ability to inhibit or enhance signal propagation through a network, when the parent gene or the protein itself is either over- or under-expressed [7].

Protein networks can be large (1000s of proteins) and contain the components of multiple known signalling pathways all joined together by other proteins whose role in the network is poorly understood. Therefore it is useful to divide PPI networks into communities (or clusters) based on their connecting architecture, under the assumption that shared network topology (interconnectedness) often correlates with shared function (or dysfunction).

Many algorithms exist to solve the problem of identifying communities in the networks, including popular Modularity-based algorithms: Fast-Greedy detection algorithm [8], Walktrap [9], SpinGlass [10], Spectral based leading-eigenvector [11] with optional fine-tuning [12] algorithms, and the Louvain algorithm [13]. Non-Modularity based algorithms include InfoMAP [14, 15], and the Bayesian based Mixed-Membership Stochastic Block model (MMSBM) which can be used to find overlapping community structure [16]. Without ground truth data to test an algorithm’s community assignment against, one often has to test the stability and robustness of clusters using a consensus based clustering approach [17, 18], measuring for example the proportion of ambiguously clustered pairs (PAC) [19].

Identifying the communities within a PPI network usually provides useful insight. Community analysis has, for example, revealed mechanistic relationships where common molecular or cellular functionality is associated with specific diseases [20-22]. To study this, gene-disease and/or gene-function associations, obtained from the *topOnto* package (*https://github.com/statbio/topOnto*) and Gene Ontology (http://geneontology.org), or experimentally studied functional group annotation data (e.g. gene-synaptic in [23, 24]) can be mapped over PPI vertices. Functional and disease enrichment of clusters can then be calculated using a suitable hypothesis driven test, i.e. the hypergeometric test, and tested against a permutation study [12] to reveal the clusters that are significantly enriched for specific annotations. This kind of analysis, for example, was used to help link together the molecular pathways in the synapse, which underpin healthy neuronal function, as well as synaptic diseases [25].

Not all proteins act similarly in propagating signals, or information, through the network. It is often assumed that proteins that interact with lots of partners have more significant impact on signal propagation or on disease mechanisms when perturbed. It is also generally found that the majority of proteins interact with just a few neighbours and thus their contribution is generally predicted to be less impactful; this was also found in scale-free networks [26]. The importance of a protein in propagating information appears to be dependent on its nearest neighbours, as well as its ability to influence other communities relative to the one it most likely belongs to. A useful network measure to quantify this is Bridgeness [27], which takes into account a vertex’s shared community membership together with its local neighbourhood. Therefore in the study of PPI networks, Bridgeness can help determine which proteins are more likely to propagate information across multiple communities simultaneously.

It was recently demonstrated that within a large-scale molecular network, the location of each sub-network of disease associated genes correlates with its pathobiological relationship to other disease sub-networks (referred to as disease modules by [28]). Diseases with overlapping modules show significant similarities at the level of gene co-expression patterns, clinical phenotype, and comorbidity. Conversely diseases residing in separated network neighbourhoods appear to be more phenotypically distinct.

Many individual software tools have been developed to address the basic steps required for the type of analysis described above. For example, the *igraph* package in R supports building a network, estimating popular centrality measures and several types of clustering [29]. Cytoscape, [30] and its plugins, support graphical representation and reconstruction of molecular networks and provide a number of approaches to extract interactions from a variety of sources such as the STRING interaction database [31], and perform clustering and estimation of the main centrality measures. Various other tools exist for functional enrichment analysis based on the GO, KEGG [32] and Reactome [33] ontologies such as the widely used DAVID package [34], GOrilla [35], BiNGO [36], and GSEA [37], to name a few. The Bioconductor [38] package *ClusterProfiler* allows estimation of the annotation overrepresentation for the clusters within a network [39] and the package *fgsea* provides fast enrichment analysis against an arbitrary set of annotations [40].

To enable rapid and systematic analysis of biological networks we designed BioNAR. The package supports a range of network analysis functions, integrating and complementing existing R packages where available, and filling the methodological gaps necessary to interrogate biomedical networks with respect to functional and disease domains. We provide a detailed topologically based network analysis pipeline, enabling the researcher to load networks generated and/or annotated using their lab’s own meta-data, thus making the tool as widely applicable and flexible as possible.

As our previous and ongoing work is related to synaptic proteome, we illustrate the package functionality with two publicly available synaptic networks (1^st^ and 2^nd^ case studies). However, BioNAR is not limited to synaptic or even proteomic networks and can be applied to any biological network, e.g., patient network, Gene-Disease (3^rd^ case study), or Disease-Disease Interactions with any customized annotation.

## Results

We developed the Bioconductor package BioNAR to support the analysis of biological networks based around a high-level pipeline presented in Figure 1. Each of these key functions is elaborated below (see also package vignette in Supplementary files for further illustration).

**Figure 1.**
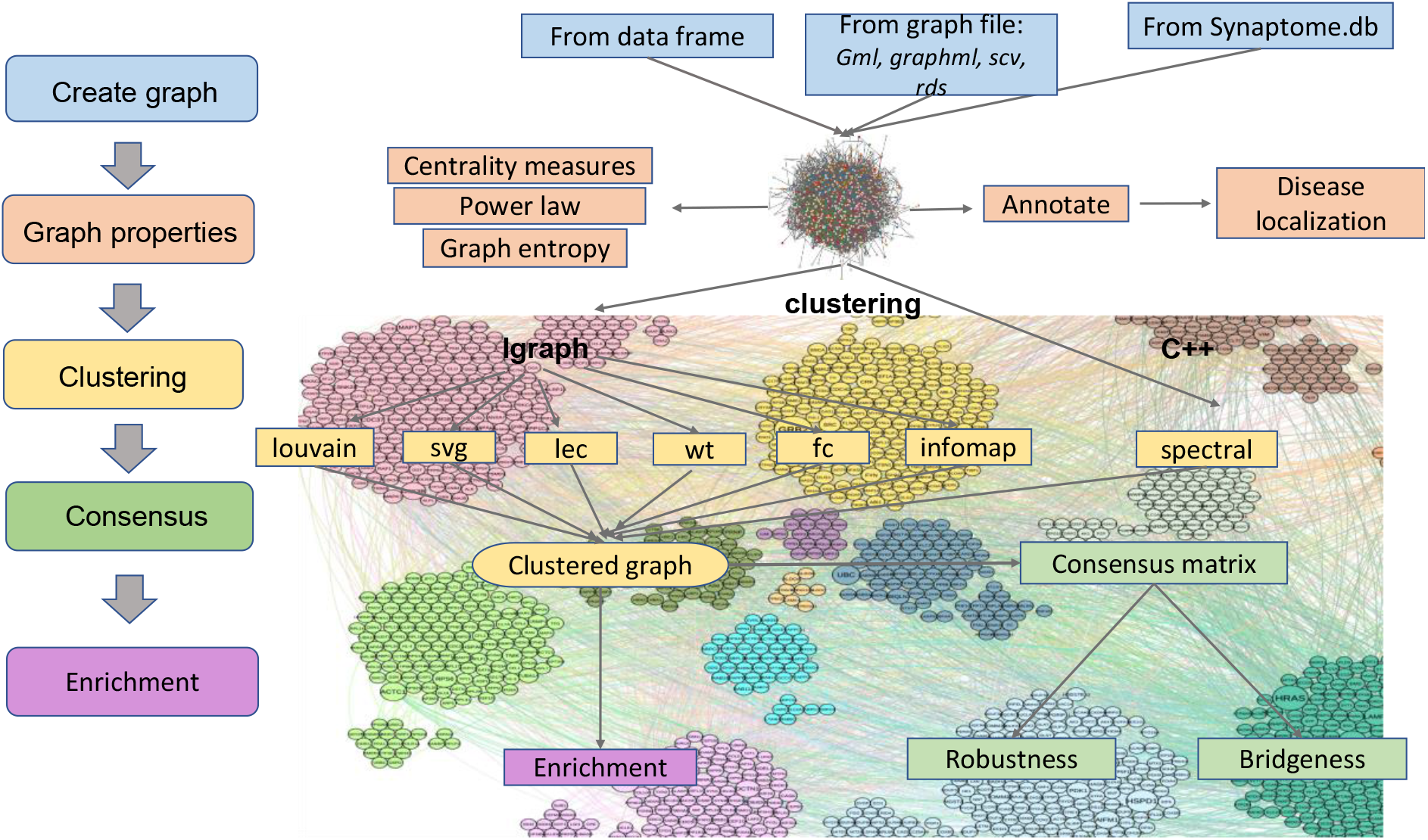
Logical outline of the network analysis pipeline implemented in the BioNAR package. Although the process is often iterative, the general flow starts with graph creation and proceeds as illustrated on the left. Each of the steps indicated corresponds to the analysis stages described in the Results section.

### Step 1. Creating a network instance

BioNAR implements networks as R data frames, where each row corresponds to a vertex interactor pair and where each vertex has a unique *vertex_ID*. Alternatively, a network can be imported from a standard graph file formats including gml, using the *igraph’*s package built-in functionality. BioNAR also allows network import for specific synaptic protein set/synaptic compartments/brain region directly from the *Synaptome*.*db* package [41]. An example of this is shown for the presynaptic case study 2. For constructing protein interaction networks described here, we selected the corresponding NCBI Gene Entrez ID to use as a unique *vertex_ID* for each node/protein.

### Step 2. Adding annotation to the vertices

Once a network is constructed, vertices are typically annotated with categorical or continuous metadata. Annotations are handled in a three-column data frame format, where the first column contains the *annotation term ID*, the second - *annotation term name*, and the third column – associated *vertex_ID*. All annotation terms for the same *vertex_ID* will be collected and stored as a string in the vertex annotation. BioNAR is designed to assign the results of any vertex calculation as a new vertex attribute, which allows intermediate results to be stored directly within the network. This supports reproducibility as many algorithms used in network analysis have a stochastic component, so each invocation can create a range of slightly different results.

In our proteomic network examples, annotations would typically include GO annotations, gene-disease association values (GDAs), gene expression values, pathway membership data and so on but they can contain any annotation data the user would be interested in. As an example of adding custom data (in case study 2), we added two new binary annotation types, one for membership of a post-synaptic cellular component (Schizophrenia- associated genes) and one for membership of a pre-synaptic compartments (presynaptic protein functional groups), using compartment-specific synaptic functional analysis reported in [23] and [42]. For disease-disease interactions or enrichment analysis such strings are converted into semicolon-separated lists to store all annotation of the vertex. For example, a protein could be annotated with two different molecular functions A and B, so in vertex annotation it will be stored as A; B.

### Step 3. Estimating network vertex properties and underlaying structure

The BioNAR package directly supports calculation of the following network vertex centrality measures, many of which are implemented in *igraph* [29]: degree (DEG), betweenness (BET), clustering coefficient (CC), semilocal centrality (SL), mean shortest path (mnSP), page rank (PR) and standard deviation of the shortest path (sdSP) (see *igraph* manual for details). Vertex centrality values can be added as vertex attributes (*calcCentrality*) or returned as an R matrix (*makeCentralityMatrix*), depending on user preference. Any other numerical characteristics, calculated for vertices and represented in a matrix form, can also be stored as a vertex attribute (*applyMatrixToGraph*).

To enable comparison of an observed network’s vertex centrality values to those of an equivalently sized randomised graph, we enabled three randomisation models including G(n,p) Erdos-Renyi model [43], Barabasi-Albert model [44], and the derivation of a new randomised graph from a given graph by iteratively and randomly adding/removing edges. To examine a network for underlying structure (i.e., not a random network), one can test a network’s degree distribution for evidence of scale-free structure and compare it against an equivalent randomised network model. For this we used the R *PoweRlaw* package (version 0.50.0) [45], which uses a goodness-of-fit approach to estimate the lower bound and the scaling parameter of the discrete power-law distribution for the optimal description of the graph degree distribution.

For proteomic networks where we also have multi-condition gene expression data, scale-free structure can also be tested by using the expression data to perform a perturbation analysis on the network to measure network entropy [7]. This kind of analysis is most useful for comparing a control relative to a perturbed network (e.g., wild-type vs cancer, untreated vs treated), where vertices with low entropy rate appear to be the most important players in disease propagation. Each protein is perturbed through over-expression and under-expression, with the global entropy rate (SR) after each protein’s perturbation being plotted against the log of the protein’s degree. In our Case Study 2, we observed a bi-modal response between over-expressed genes and their degree and opposing bi-phasic response relative to gene under-expression and degree (Figure 3C). This type of bi-modal, bi-phasic behaviour has been observed only in networks with scale-free or approximate scale-free topology [7].

### Step 4: Clustering

BioNAR supports a non-exhaustive set of commonly used clustering algorithms. These are Modularity-Maximisation based algorithms, including the popular agglomerative ‘Fast-Greedy Community’ algorithm (fc) [8], process driven agglomerative random walk algorithm ‘Walktrap’ (wt) [9], and coupled Potts/Simulated Annealing algorithm ‘SpinGlass’ (sg) [10, 46], the divisive spectral based ‘Leading-Eigenvector’ (lec) [11] and fine-tuning (Spectral) [12] algorithms, and the hierarchical agglomerative ‘Louvain’ algorithm (louvain) [13]. We also included a non-Modularity information-theory based algorithm ‘InfoMAP’ (infomap) [14, 15].

All algorithm implementations, apart from Spectral, were imported from R’s *igraph* package [29]. The Spectral algorithm [12] was written in C++ and wrapped in R within a satellite CRAN package *rSpectral*, (https://cran.r-project.org/web/packages/rSpectral/index.html), linked to BioNAR (More details in Supplementary Methods).

Depending on the purpose of the study all clustering algorithms can be applied to the network under investigation simultaneously, with each algorithm’s community membership stored as a vertex attribute. The user also has the option to select specific clustering algorithms to run over their network, since running all clustering algorithms over the large network might be time consuming. For instance, it took approximately half an hour to run all nine algorithms on the presynaptic network described in Case Study 2 on a modern MacBookPro laptop.

A common phenomenon when applying Modularity based clustering algorithms over networks of a large size, is to end with large, or ‘super’, communities which masks the networks substructure. In this situation we provide the user the facility to re-cluster these large/super communities in a hierarchical manner, applying the same, or potentially a differing, clustering algorithm at each iteration (*recluster*).

To compare the usefulness of different clustering algorithms on a network, a summary matrix can be created, consisting of: the maximum Modularity obtained (mod), the number of detected communities (C), the number of singlet communities (Cn1), the number of communities with size >= 100 (Cn100), the fraction of edges lying between communities (mu), the size of the smallest community (Min. C) and the largest community (Max. C), the average (Mean C), median (Median C), first quartile (1st Qu. C), and third quartile (3rd Qu. C) of community size (Table 1).

**Table 1.**
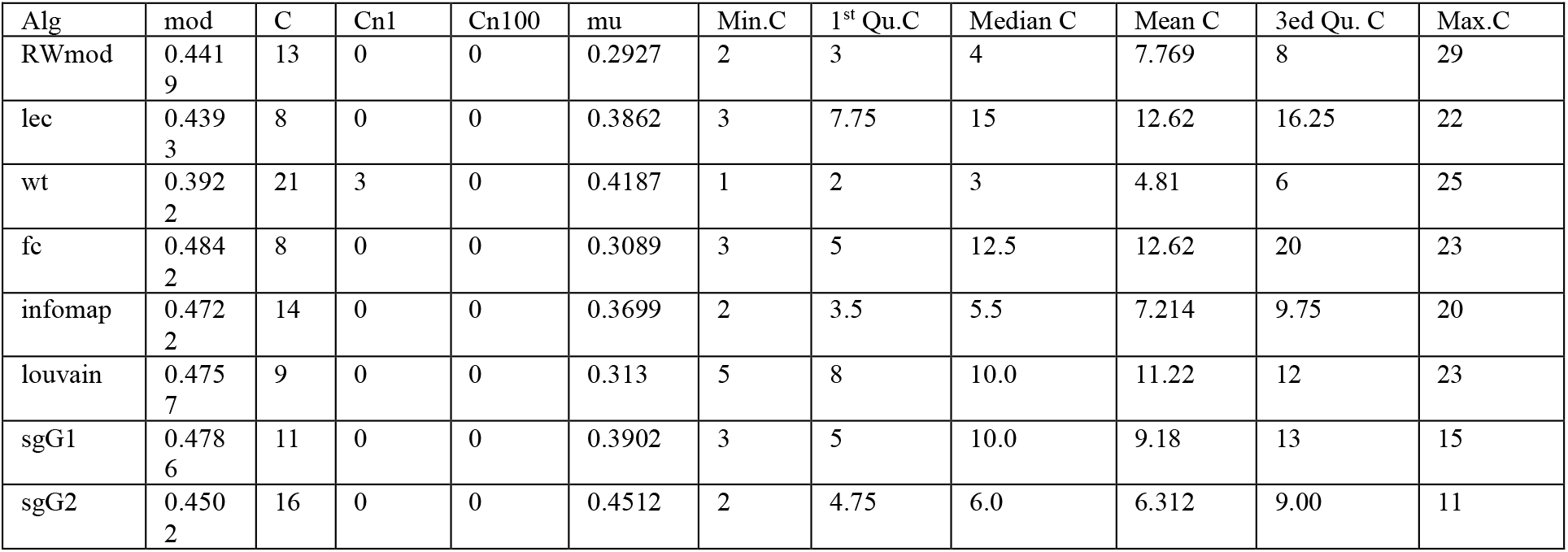
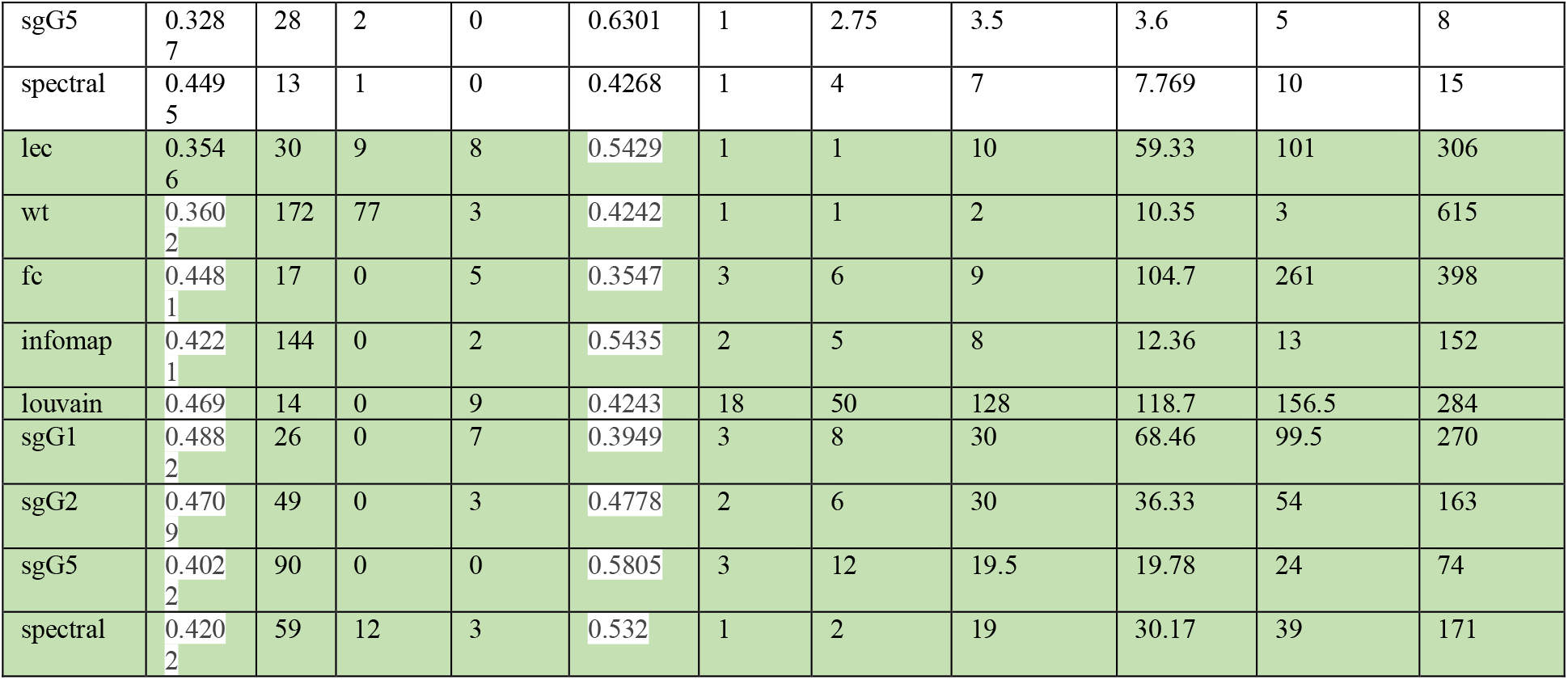
Clustering summary for MASC (upper white) and presynaptic (lower green) networks. Shown are the maximum modularity obtained (mod), the number of detected communities (C), the number of singlet communities (Cn1), the number of communities with size >= 100 (Cn100), the fraction of edges lying between communities (mu), the size of the smallest community (Min. C) and the largest community (Max. C), the average (Mean C), median (Median C), first quartile (1st Qu. C), and third quartile (3rd Qu. C) of community size.

To test the robustness of communities found by a clustering algorithm, a consensus matrix is built by randomly selecting a proportion (by default 80%) of the network vertices and rerunning the clustering algorithm (by default set to 500 times). The functionality provided in the R package *clusterCons* [18] is then applied to test the robustness of discovered communities, where community robustness values range from 0, indicating no confidence in the community existing, to 1, indicating absolute confidence in the cluster existing.

The BioNAR package provides functionality to visualise a network’s community structure with our implementation of cluster-driven layout, which is suitable for even the largest network (i.e., tens of thousands of vertices and millions of edges). This layout splits the network into clusters, lays out each cluster individually, and then combines individual layouts with the *igraph* function merge_coords, so that each distinct community is shown independently and painted in a unique colour.

To allow comparison of networks with different structures, we implemented a normalised modularity measure [47-50], which is defined as:

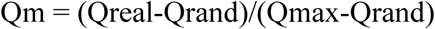

Where Qreal is the network modularity of a real-world signaling network and, Qrand is the average network modularity value obtained from 10,000 randomised networks constructed from its real-world network. Qmax was estimated as: 1 − 1/M, where M is the number of modules in the real network. Here, randomized networks were generated from a real-world network using the edge-rewiring algorithm [51].

### Step 5. Bridgeness and identifying “influential” vertices

The clustering algorithms we used in Step 5 place each vertex into a single cluster, which in many cases is an oversimplification. The bridgeness metric measures the probability that a vertex belongs to more than one community at the same time. bridgeness can be estimated from the consensus matrix calculated above, taking values between 0 - implying a vertex clearly belongs in a single community, to 1 - implying a vertex forms a ‘global bridge’ across every community with the same strength [27, 52]. More details in Supplementary Methods.

Although useful itself, bridgeness becomes especially informative when combined with other vertex centrality measures, e.g., semi-local centrality, which considers the nearest and next to the nearest vertex neighbours. It also lies between 0 and 1 indicating whether the vertex is likely to have local influence.

Plotting bridgeness against semi-local centrality, allows us to categorise the local and global influence of each vertex within a network given only the network structure (see case study 2). BioNAR also supports the comparison of bridgeness against any vertex centrality measure (or any normalised numeric vertex value) of the user’s choice, e.g., against Page Rank.

In the context of proteomic networks, we know that proteins are often present in multiple copies, in multiple subcomplexes sometimes serving very different biological functions. Thus, estimating the bridgeness allows each protein in the network to be ranked according to its predicted importance for propagating signals through the network based on architecture alone.

### Step 6: Studying the overlap or separation of annotation pairs

Given two annotations that are distributed across a network, a common query is to find the points of intersection where the two annotation sets overlap (or segregate). To support such queries, we implemented the algorithm from [28], which tests if the observed mean shortest paths between two distinct annotation sets, superimposed on a network, is significant compared to a randomly annotated network (More details in Supplementary Methods).

This method is often applied to disease annotations although any similar type of annotation will work. The BioNAR command *calcDiseasePairs* calculates the observed overlap between two annotation sets on a network, and compares this to a single instance of the network with annotations randomly permuted; this is useful for a quick estimate of how likely the overlap is simply a random occurrence.

To calculate the significance of observed overlaps (or separations) the observed annotation pairs on the network the command *runPermDisease* should be used. This compares the overlap against multiple permutations of the network (where the user can define the number of permutations). Executing this command, which may take time depending on the number of permutations chosen, generates a results table containing the overlap of each annotation pair with p-value, p.adjusted by Bonferroni test, and q-value.

### Step 7: Enrichment Analysis

Over-representation analysis (ORA) is a common approach to identify annotation terms that are significantly over- or under-represented in a given set of vertices compared to a random distribution.

In biological networks GeneOntology terms and Pathway names are amongst the most frequently used. ORA differs from Gene Set Enrichment Analysis (GSEA) as the latter use numerical values associated with genes, such as expression value, while the former relies on null hypothesis tests, such as the hypergeometric test statistics. The most commonly used GeneOntology analysis can be performed with dedicated Bioconductor tools, such as *clusterProfiler*. To keep the package as general purpose as possible, we used the Bioconductor package *fgsea* to implement ORA functionality on top of arbitrary string vertex annotation and vertex grouping, obtained for example, by clustering. We represent the results of ORA as a R data frame, with rows representing the group of vertices and columns the p.value enrichment values for set of annotations terms under study. We also provide p-value, adjusted p-value, size of overlap and list of vertices that contribute to the annotation term.

We now outline three brief use-cases. The supporting files and documentation/video walkthroughs are available as supplementary materials and provide an entry point to anyone wanting to use these as starting points for similar analysis.

### Case study 1. MASC network

The first test case chosen is a previously studied NMDA receptor complex known as the MASC network [53]. This is a relatively small PPI network, representing a protein complex surrounding the mammalian NMDA receptors and consists of 101 proteins with 246 interactions [53]. It is a good example of the type of network that would come from a typical pull-down experiment. The original MASC study provided an analysis of the organisation and underlying functionality of the modularised MASC complex, and was clustered using the Random Walk algorithm [54] as shown in Figure 2, central. The analyses used in the original study [48] predated most of the tools listed here and were achieved using almost entirely manually curated datasets and bespoke code developed for the specific study. Using BioNAR we replicated the cluster analysis using our set of algorithms and visualised the results using Gephi (www.gephi.org). Clustering summary is presented in Table 1 (top half), which demonstrates that all algorithms give similar clustering structure with 8-13 communities, composed of an average of 5-10 protein components each.

**Figure 2.**
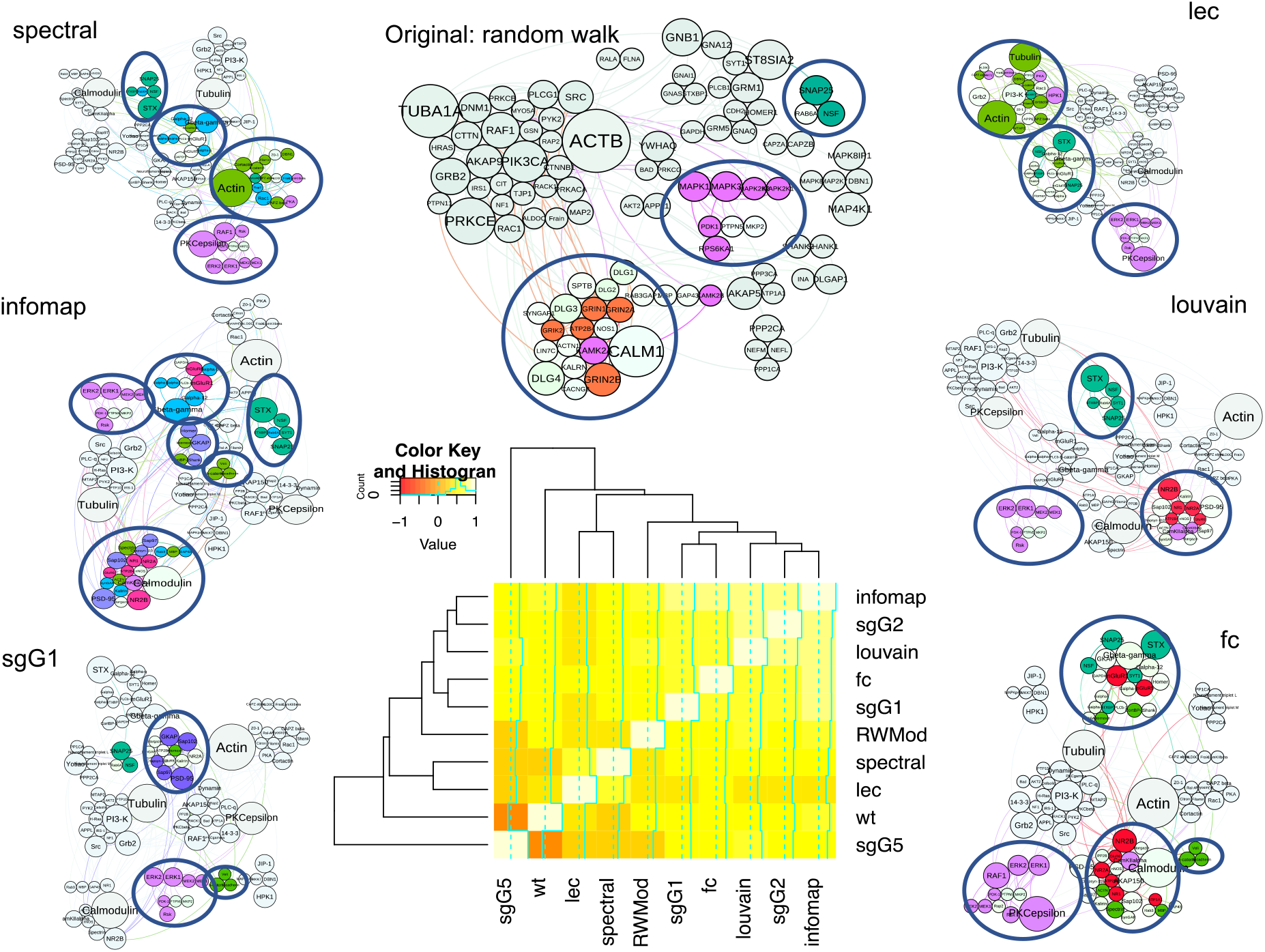
Shown the clustering results for Case Study 1 (MASC network) for 6 algorithms. The colour code corresponds to the protein functional families described in the original paper [53] as follows: red – channels and receptors, light green – cell adhesion and cytoskeleton, dark green – synaptic vesicles/ protein transport, light blue – G-proteins and modulators, purple- MAGUKs/adaptors/scaffolds, maroon – kinases. Highlighted are only the clusters with significant enrichment (p-val < 0.05) with specific family. Figure in middle bottom illustrates the similarity of algorithm’s performance by Reduced Mutual Information (RMI) index (similar algorithms are coloured yellow, different – red).

We performed overrepresentation analysis to test if the obtained community structures (clusters) have meaningful enrichment for functional annotation terms, using the functional protein family annotations from the original paper [53]. As can be seen from Figure 2, all clustering algorithms split the network into similar functional units. With all considered algorithms we find two persistent clusters, containing proteins associated with: 1) Synaptic vesicles/protein transport, containing 2-5 proteins of 6 possible family members, and 2) kinases, containing 4-7 from 18 family proteins. In addition, the following families appear enriched in the clusters produced by the majority of algorithms, e.g., 1) channels and receptors, containing 5-6 proteins of 8 family members (RWMod, Louvain, fc, infomap, sgG2 algorithms), 2) cell adhesion and cytoskeleton, containing 3-7 proteins from the respective family (spectral, lec, wt, fc, infomap, sgG1), 3) G-proteins and modulators, containing 3-5 from 17 possible proteins (spectral, infomap, sgG2, sgG5), 4) MAGUKs/adaptors/scaffolds (3-5 from 12 family proteins, identified by wt, infomap, sgG1, sgG2). P-value and p.adj values for the respective algorithms can be found in Supplementary table 1. In terms of overrepresentation, the highest levels of significant enrichment is observed in the clustering produced by the infomap algorithm, which provides the split into 5 distinct significantly enriched functional communities (Figure 2, Supplementary Table 1).

We compared the performance of different clustering algorithms by estimating Newman’s Reduced Mutual Information (RMI) index [55], which enables pairwise comparison of classifications of the same sets of objects. infomap, Louvain, fc, sgG1, sgG2 algorithms split the network in similar way to the original clustering, while lec, spectral and, especially, wt give more divergent results (Figure 2 bottom).

### Case study 2. Presynaptic network

A scale up, in both size and complexity, from the MASC network in study 1 is the study of an entire subcellular compartment’s protein-protein interaction network that integrates data from multiple experiments. For that we constructed a proteome network for the entire presynaptic compartment using the R package Synaptome.db [41]: 2304 vertices were extracted from published studies of this compartment, which were combined with protein-protein interactions from Synaptome.db to obtain a Largest Connected Component containing 1780 vertices connected by 6620 edges.

We examined the network for underlying structure (i.e., not random) by testing the network’s degree distribution against two randomised network models, and found evidence of scale-free structure in the network: an alpha (goodness-of-fit) score to a power-law distribution of 0.25, was calculated using the *PowerLaw* function (Figure 3C, left panel). Supporting evidence for scale-free structure was obtained by performing a perturbation analysis on the network [7]. As in [3], proteins were set to initial values of 2 with perturbed values of 14 when modelling activity and set to initial values of 16 with perturbed values of – 14 when modelling inactivity. Then each protein was perturbed through over-expression (Figure 3C, red) and under-expression (Figure 3C, green), with the global graph entropy rate (SR) after each protein perturbation being plotted against the log of the protein’s degree (Figure C3, right panel). We observed a bi-modal response, between gene over-expression and degree, and opposing bi-phasic response relative to over/under-expression between global graph entropy rate and degree. This type of bi-modal, bi-phasic behaviour has been observed only in networks with scale-free or approximate scale-free topology [7].

**Figure 3.**
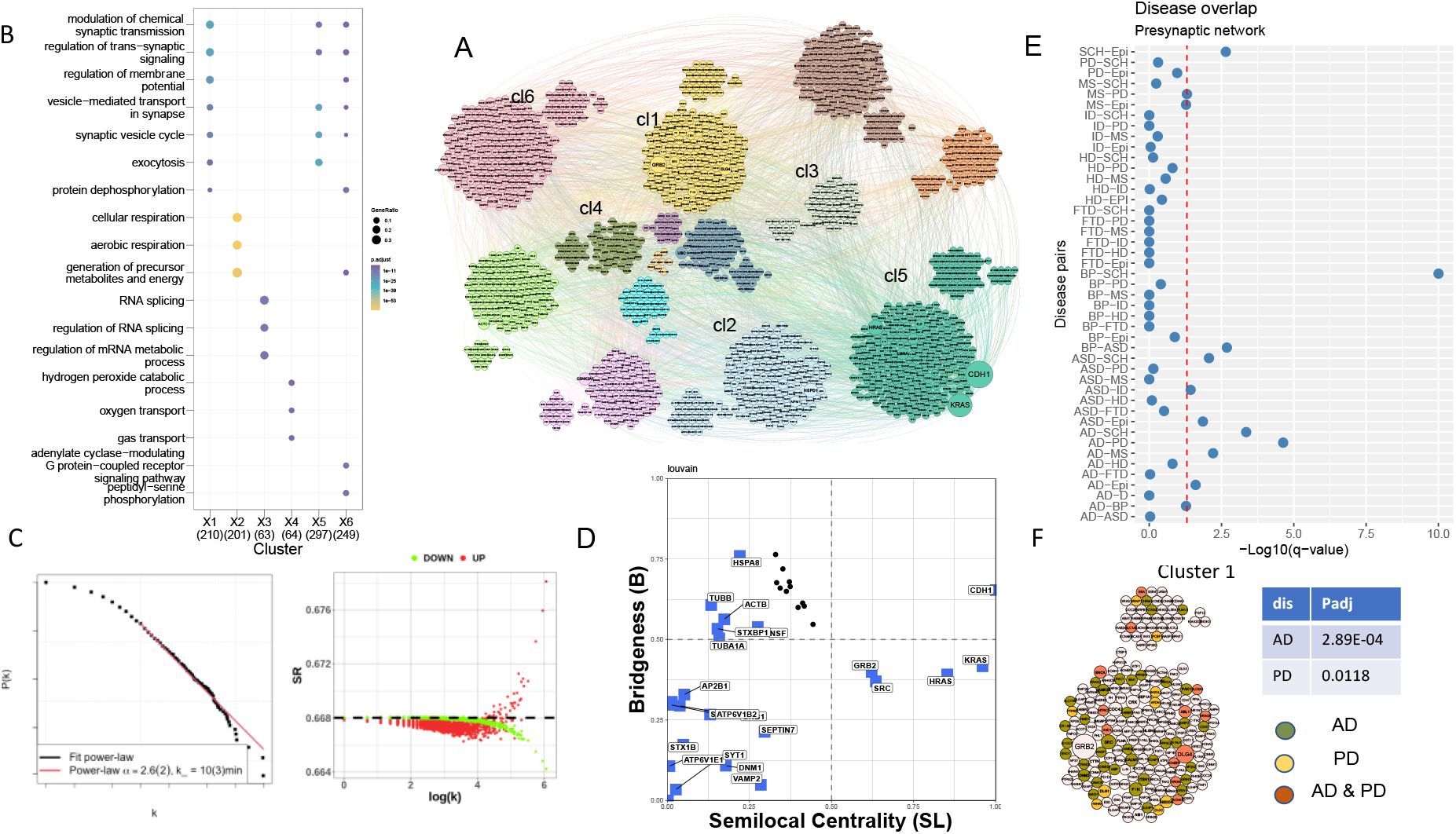
Presynaptic network, analysis results. A. clustering results from applying the Louvain algorithm, with clusters assigned a unique colour. B. Enrichment analysis results performed using the ClusterProfiler package for the 1-6 communities (highlighted at Figure 3A). C. Power law fit - shown the log-log plot of the CDF of presynaptic PPI network degree distribution (P(k)), versus its degree (k), with the best fitting power-law distribution to the network data highlighted in red. D. Bridgeness results shown for louvain algorithm, highlighted are the genes most frequently found in presynaptic compartment. E. Disease- disease overlap for presynaptic compartment, red dotted line shows the confidence cut-off (q val <0.05). F. Cluster 1 in details with highlighted proteins associated with AD and PD.

The presynaptic PPI network was also analysed with respect to vertex centrality measures including, Degree (DEG), Betweenness (Bet), Closeness, Clustering Coefficent (CC), Page Rank (PR), Semi-Local centrality (SL), and mean shortest path (SP) with the results presented in Supplementary Table 2.

To detect community structure in the presynaptic PPI network we used a set of commonly used clustering algorithms implemented in BioNAR. A summary of the clustering results from each algorithm applied to the presynaptic network is presented in Table 1 (green bottom). As seen in Table 1 each algorithm produces a different number, and size, of communities. For illustration purposes we selected clustering results from “Louvain” algorithm, which gives a reasonably small number of functionally enriched communities (14), no singletons, and communities distributed in size from 18 (the smallest) to 284 (the largest) vertices (Figure 3A) for the further analysis:

A snapshot of results for enrichment for Gene Ontology Biological Process terms, using the ClusterProfiler package [39] is shown in Figure 3B, where we highlighted the first 6 most enriched clusters (full enrichment results are given in Supplementary Table 2). Enrichment for GO functions are clearly distributed across different communities. For example, “regulation of trans-synaptic signalling”, “regulation of membrane potential”, “regulation of chemical synaptic transmission” terms are associated with cluster 1, synaptic vesicle cycle- related terms – with cluster 5, RNA splicing and metabolic processes - with Cluster 3, and “oxygen transport” and “hydrogen peroxide catabolic process” within Cluster 4.

To test specific synaptic functions associated with clusters we additionally annotated our graph with the SynGo annotation [56](release 20210225) and schizophrenia-related annotation (SHAnno) presented in [23], and performed over-representation analysis on the Louvain clustered presynaptic network, using all three sets of annotations. The full cluster overrepresentation results are presented in Supplementary Table 3, but briefly, in agreement with results from ClusterProfiler; cluster 1 is overrepresented with “structural constituent of postsynaptic density” (p.adj = 5.18E-06), “regulation of postsynaptic neurotransmitter receptor activity” (p.adj = 7.27E-03) and “excitability” (p.adj = 7.64E-4) terms, which may indicate that cluster 1 is enriched with membrane associated proteins from both the pre- and post-synaptic sides; it is possible that some proteins may be contaminated from the post- synaptic membrane, but the majority have been annotated elsewhere as being both pre- and post-synaptic (for example by SynGo).

From a core set of five clustering algorithms including infomap, Spectral, sgG1, Louvain and fc algorithms (Supplementary Table 4), we identified 324 (324/1780 ∼18%) potential bridging proteins (Bridgeness value >= 0.5) within the presynaptic PPI network; 15 of these were identified by all five clustering algorithms (15/323, 4%), 109 by three and more (109/323, 33%). The bridging proteins were found distributed through the clustered network, and include for example, STXBP1, ACTN1, CDH1, APP, VCP, PTPRF CAMK2A, CAMK2B; many of these proteins are understood to be important in forming cytoplasmic scaffolds that organize and connect the synaptic vesicle with the presynaptic membrane or are involved in multiple signalling cascades [42].

We checked if any bridging proteins were also annotated with one or more of the more common synaptopathies, specifically: Alzheimer disease (AD), Bipolar Disorder (BP), Autistic spectral disorder (ASD), Epilepsy (Epi), Parkinson disease (PD), Schizophrenia (SCH), Frontotemporal dementia (FTD), Intellectual Disability (ID). Of the 324 proteins, 169 (169/324∼51%, p = 8.56E-05) were annotated with at least one disease. Of 15 proteins bridging proteins found in all five clustering algorithms, 10 (10/15 ∼67%) were found associated with at least one synaptic disease given in our set (Supplementary Table 4).

The plot of bridgeness, using the Louvain clustering algorithm, against semi-Local centrality is shown in Figure 3D. We highlighted 19 bridging proteins found most frequently in the presynaptic compartment (found in 15 and more presynaptic studies [41]). Among these, six have high bridgeness values, thus likely have a global influence over network: HSPA8 (AD, PD, SCH), ACTB (PD, SCH), TUBB (FTD, SCH), STXBP1(AD, PD, FTD, SCH, ID), NSF (PD), TUBA1A (SCH). Additionally, we highlighted CDH1, as it has both high values for bridgeness and semi-local centrality and is associated with ID. High local centrality values were also observed for GRB2, HRAS, NRAS, which are recognised as local hubs participating in many signalling cascades (also known as party hubs [23]).

Many neurological disorders are co-morbid and share similarities in clinical phenotype. To test whether common synaptic molecular mechanisms might underpin these disease similarities we performed the disease separation analysis for the disease annotations set, mentioned above (Figure 2E, more detail in Methods). The most overlapped pair was BD - SCH (q.val = 1.19E-09), followed by AD-PD (q.val = 1.64E-05), AD-SCH (q.val = 4.12E- 04), BP-ASD (q.val = 7.2E-04) (Supplementary Table 5), which are already known for their comorbidity. From the analysis of distribution of disease associated proteins over the network combined with overrepresentation analysis, it appears that the majority of diseases, for instance, AD, PD, EPI, ASD, SCH, ID and even BP are overrepresented in cluster1, and are associated with proteins annotated with major synaptic functions like “trans-synaptic signaling”, “synaptic vesicle cycle”, etc (Figure 1F, Supplementary Table 3, highlighted in red). The co-occurrence of enrichment for specific synaptic functions and disease associations in the network clusters points to shared molecular mechanism.

This case study provides an illustration of how proteomics datasets from diverse studies can be integrated into a single large model. BioNAR then provides a suite of analytical tools that can extract key structure/functional relationships within the data than can even provide a novel perspectives on the interplay between the molecular mechanisms of different diseases.

### Case study 3. Human Disease Network

In 2007 Barabasi and colleagues curated and published a Human ‘diseasome’ network along with an analysis of two of its natural projections: Human Disease Network (HDN) and Disease Gene Network (DGN) [57]. The diseasome was built by collecting and annotating relationships between known human diseases, and disease-causing gene mutations. The diseasesome is a bipartite network as it only contains the edges between two vertex types, i.e., ‘disease’ and ‘gene’ vertices. Projections of a bipartite network result in mono-type graphs, where a pair of vertices are connected if, and only if, that pair was connected to the same, but opposite, typed vertex in the original bipartite network. Therefore, the HDN graph contains disease vertices and connections between them if both diseases are linked by mutations in the same gene, while the DGN graph contains only gene vertices, which are connected only if both genes are associated with the same disease. We have reconstructed all three networks from the supplementary material in[57]. All three networks, i.e. diseasome, HDN and DGN contain a largest connected component (LCC), containing approximately 46%, 40% and 50% of all possible vertices, respectively.

When we randomly permuted connections in the original diseasome bipartite network and compared the size of the LCC in all three networks, we found each to be significantly smaller (pval < 0.002) compared to that in the unperturbed networks (See Figure 4, A). It was discussed in [57], that the small size of the LCC was probably caused by preferential attachment of genes within the same same disorder class.

**Figure 4.**
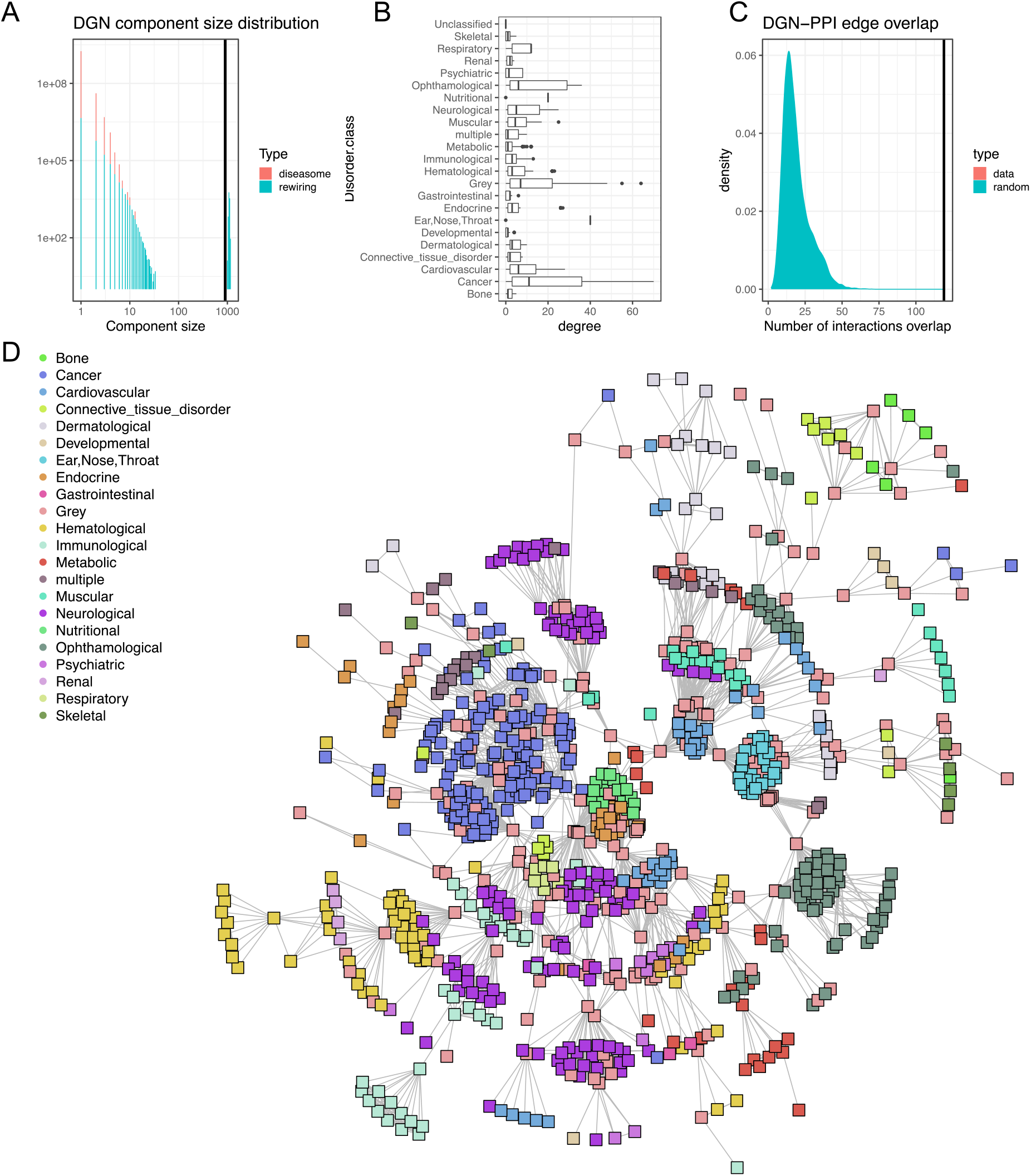
Disease Gene Network (DGN) analysis performed with BioNAR A. Distribution of connected component sizes in DGN. Red columns correspond to component sizes in the original network, while green to randomly perturbed networks. Vertical line on the graph corresponds to the size of the giant component of the original DGN, which includes 50.8% of nodes. Like in the original Barabasi paper, its size (903) is significantly lower (pval<10^−3^) than the average size of the giant component (1088±21) in the set of rewired networks. B Degree distribution of genes annotated with different disorder classes. It can be seen that high-degree genes come almost exclusively from the disorder class Cancer. A further disorder class ‘Grey’, which also contains high degree genes, is associated with many disorders [57]. C. Number of observed interactions between genes annotated by the same disorder type (vertical line) and distribution of the expected numbers from randomized network. D. The largest connected component of DGN. Nodes are coloured according to the disorder class.

Here we focussed on the analysis of DGN (Figure 4, D); the equivalent HDN analysis is available in Supplementary material. Unlike the networks in the other case studies, analysed in this paper, the DGN network does not show a scale-free distribution (Supplementary Figure 2).

The degree distribution for the DGN network only follows a power-law at the extreme high degree tail (Supp. Figure 2). This can be attributed to the DGN’s construction since we project an incomplete bipartite disease-gene network onto gene nodes. The low-degree genes in such a network result from our sparse knowledge about ‘less studied’ diseases, while the well-studied diseases such as cancers, with high coverage in genes, lead to high degree and well-established connectivity of vertices (Figure 4, B), which reflects in power-law-like behaviour in this range of degrees.

Similar to [57] we compared the inferred interactions between genes in GDN with experimentally observed ones. While [57] used a manually curated PPI network for DGN, we extracted a PPI from published synaptic proteome data available via the synaptome.db Bioconductor package [41] by filtering out vertices from the ‘neuroPPI’ network, which were not found in DGN. We calculated the number of disease module interactions as the number of PPI edges found between genes annotated to the disorder class given in DGN (Fig 4, C). We observed a smaller number of disease module interactions (119), than that found in [57] (290) (which is not surprising given we are focussing on just synaptic interactions), but this was still significantly larger than expected by chance (19±9.1) (Figure 4,C), giving indication that genes associated with the same disease are highly likely to physically interact.

Thus, we reproduced a major part of the [57] analysis on human diseasome in the context of a network derived from primary synaptic data within BioNAR framework which demonstrates how integration of these different Bioconductor packages in BioNAR can enable deeper understanding of the biological system at hand.

The code and datasets for three cases are available from https://github.com/lptolik/BioNAR_paper_supplementary and https://datashare.ed.ac.uk/handle/10283/4793

## Conclusion

BioNAR provides a pipeline that implements many core network analysis techniques, including network import and annotation, estimating scale-freeness and a range of centrality measures, and providing nine different clustering algorithms. It is further extended with methods to estimate network entropy and normalized modularity that can be used to compare networks with different structures. The implementation of bridgeness can be used to estimate the likely importance proteins may have in propagation of signals between different communities. Annotation enrichment features can be used to identify co-ocurrence of annotation pairs (typically disease) within a network. Beyond this, the package also allows users to estimate the robustness of a clustered through cross-validation against a selection of randomized network structures to evaluate the significance.

It can be seen from our synaptic network case studies that for smaller networks like the MASC complex, clustering results are mostly similar to each other (Table 1). However, as the network’s size increases, like in the presynaptic network example in Case Study 2, clustering results produced by different algorithms vary more, so the user will have to select a preferred clustering method based on the clustering summary provided (number of communities, their sizes, etc). Enrichment results for functional annotations in the clusters provide a potential test for the best clustering for the biological network of interest but the approach relies on an assumption that functional annotation is sufficiently complete and accurate. Enrichment analysis can be performed either with standard ontologies such as GO or HDO, or with an in-house ontology or annotation(s) that can be assigned to the network components.

Proteins can belong to different molecular complexes simultaneously either spatially due to multiple expressed copies of the same molecule or temporally as a result of post-translational modifications occurring in signalling cascades. Therefore, we enabled a function that takes into account the probability of the vertex to belong to different communities, under assumption that more communities the vertex can belong to, the more influence it has on network topology and signalling. We consider ‘influential’ or ‘bridging’ proteins to be those known to interact with many neighbours simultaneously, helping organise function inside communities (Han et al., 2004) they belong to, but that also affect/influence other communities. In our presynaptic network (case study 2), we found the majority of identified bridging proteins were associated with one or more synaptopathies, thus supporting the hypothesis that these proteins may play important roles in function and disease.

Methods embedded in BioNAR can be used to extract a molecular signature for a disease, or function, at the network level. In the case study 2, the presynaptic network, we found a few neurological disorders to overlap with one another with a high level of significance, e.g. Alzheimer Disease and Parkinson Disease, Schizophrenia and Bipolar Disorder, Autism and Intellectual Disability, and, indeed, their comorbidity is confirmed in the literature [58-60]. The same analysis can be performed with any in-house annotation using BioNAR functionality, which may give new and unexpected relationships between different diseases or functions.

For this first release of the BioNAR package we tried to include the most commonly used methods of network analysis. From a technical perspective, BioNAR could be extended to improve support for high performance compute clusters for analysis of very large networks; integration with graphical databases such as Neo4J for more efficient storage, retrieval and querying of networks and the use of network embeddings, which is highly active area of research in the AI community[61-63]. Future releases of BioNAR will also see the inclusion of probabilistic network algorithms based on Bayesian inference to utilise both vertex and edge meta-data, including a probabilistic clustering algorithm built from Stochastic Block Models (SBMs) and belief propagation, to model the influence of vertex meta-data on network clustering (https://github.com/cmclean5/rblock), and network reconstruction, i.e. estimating the true network structure and quantifying its uncertainty, given multimodal edge meta-data [64] (https://github.com/cmclean5/NetworkReco); [65]. However, this first release of our package already provides researchers with a useful network analysis ‘toolkit’, integrating many of the basic network analysis tasks needed by the wider bioinformatics communities.

## Key Points

- BioNAR integrates together the most fundamental down-stream network analysis pipelines (workflows) needed in the study of biomolecular data into a single package, with an emphasis on proteomic data.
- Case Studies provide walk-through examples of proteomics analyses at both smaller focussed scale as well as much larger meta-level models produced from integrating across multiple published studies.
- BioNAR is agnostic to the type of network data, the network analysis pipelines implemented within BioNAR are applicable to the vast spectrum of biomedical networks, from patient similarity networks, disease networks, down to the family of omic networks.

## Funding

This research has received funding from the European Union’s Horizon 2020 Framework Programme for Research and Innovation [No 945539 (Human Brain Project SGA3)], and Cancer Research UK [No CTRQQR-2021\100006 (Cancer Research UK Scotland Centre)].

## Notes

https://datashare.ed.ac.uk/handle/10283/4793

### Competing Interest Statement

The authors have declared no competing interest.

